# Bred for affection: The canine anterior ectosylvian gyrus responds selectively to social reinforcement

**DOI:** 10.1101/2024.06.10.598283

**Authors:** Kai J. Miller, Frederik Lampert, Filip Mivalt, Inyong Kim, Nuri Ince, Jiwon Kim, Vaclav Kremen, Matthew R. Baker, Max A. Van den Boom, Dora Hermes, Volker A. Coenen, Gerwin Schalk, Peter Brunner, Gregory A. Worrell

## Abstract

Studying mammalian brain function aids our understanding of human brain evolution. We implanted a beagle with a prototype human neuromodulation platform that measures activity from the brain surface. One year later, a set of simple sensory tasks was performed, finding visual and somatosensory representation in the canine homologs of the expected areas in humans. Surprisingly, the canine anterior ectosylvian gyrus, which is anatomically homologous to human receptive speech areas, was selectively active during independent social reinforcement tasks. This suggests that human speech understanding may have evolved from more general mammalian brain structures that are specialized for social reinforcement.

## MAIN TEXT

Domestic dogs (Canis familiaris) have been bred for social reinforcement from humans, at first passively through domestication of the gray wolf over tens of thousands of years^1^ and then actively as cultivated breeds over the last several hundred years^2^. While all mammals are primed to social feedback from their own species, subtle changes to the canine genetic code throughout these millennia of domestication have shaped dogs’ brains to be uniquely receptive to verbal and tactile (petting) reinforcement from their human companions^3^. This unique receptiveness to humans enabled an unexpected discovery we made about how the canine brain is organized, during feasibility testing of an implanted software-hardware neuromodulation platform^4^.

We implanted a 2-year old beagle canine, “Belka”, with a 32-channel electrocorticography (ECoG) construct consisting of 3 arrays overlying the right hemisphere (Fig. 1). In the year after implant, Belka maintained a healthy social life, playing with our team daily and interacting with the pack she is housed with. Regular systems diagnostics were measured, and a set of controlled sensory tasks were performed at 11-12 months to assess viability of the platform.

**Figure 1.**
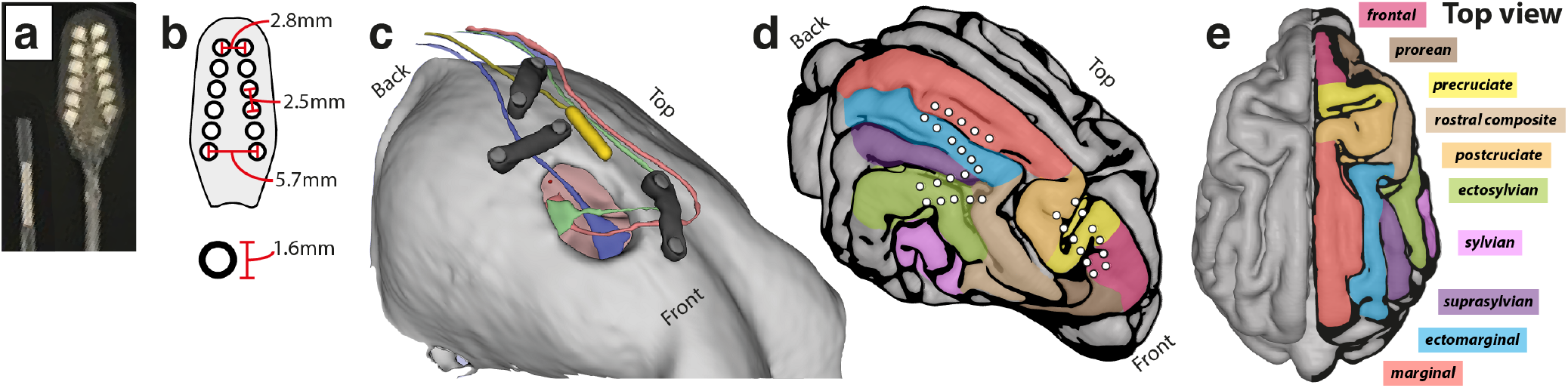
Right hemisphere implant and anatomic segmentation of the dog’s brain. **a**, The ground electrode and one of the 2×6 angled electrode arrays used for recording. **b**, Schematic with dimensions of electrode array. **c**, Rendering of skull and craniotomy showing electrode arrays in situ, and the ground electrode in yellow. Plates that hold lead in place are shown in black. **d**, Brain rendering showing three grids in situ, extracted from pre-implant MRI and registration to post-implant CT, with gyri color-coded. **e**, Top view of the canine cortex, with gyri and labels color-coded.

As with human ECoG experiments^7^, robust task-associated changes are found in the power spectral density (PSD) of the canine brain surface electrical potential: When a region is active, we measure spatially-focal broadband power increases, and, in some cases, power decreases in narrowband low-frequency oscillations (rhythms) with peaks below ∼50Hz (Fig. 2). For example, a simple lights ON-vs-OFF task reveals a robust ∼15Hz rhythm that emerges in the dark (an electrophysiological homolog of the human 8-10Hz *α*-rhythm^8^), centered over the marginal & ectomarginal gyri (Figs. 2a and Extended Data Fig. 1). A smaller region - within just the marginal gyrus - that shows broadband power increase when the lights are turned on. The visually-associated broadband increases from the marginal gyrus align with the known primary visual area of the canine brain identified from lesioning^9^ & fMRI^10^ studies. More generally, a full set of simple visual, auditory, and tactile sensory tasks demonstrate that, as with human ECoG^5,6^, local brain activity can be captured across a range of behaviors and regions by broadband spectral change from the surface of the canine brain (here captured at 65-150Hz, Fig. 2).

**Figure 2.**
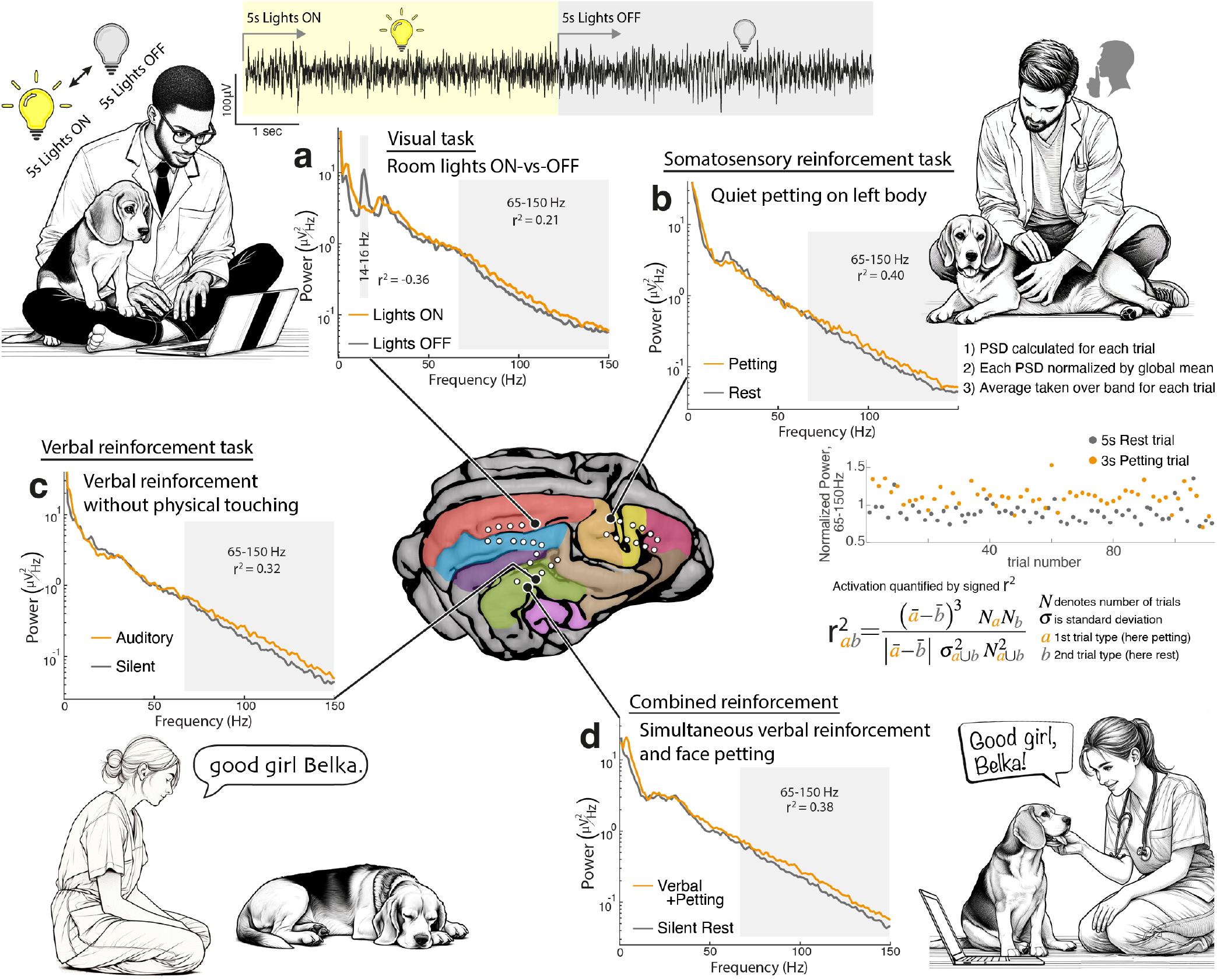
Brain electrophysiology during sensory tasks. Power spectral densities (PSDs) are shown for four different sensory tasks, comparing active (orange) and inactive (gray) behavioral states. **a, Visual task** - Blocks of 5 seconds with the room lights on (“Lights ON”) were interleaved with 5 seconds in the dark (“Lights OFF”). Signal changes are visually apparent in the raw voltage trace from the marginal gyrus, both in low frequency oscillation ranges (here at 14-16Hz) and also in broadband spectral changes (here captured at 65-150Hz), which have been shown in humans to correlate with local neuronal population activity^5,6^. **b, Somatosensory reinforcement task** - 3s blocks of tactile stimulation (petting left whiskers, front and hind limbs, and torso) were interleaved with 5s blocks of rest, with no verbal input. PSD from the post-cruciate gyrus is shown. **c, Verbal reinforcement task** - 3s blocks of verbal reinforcement, saying “Good girl Belka” once each block, were interleaved with 5s quiet blocks. There was no physical contact. PSD is shown from the anterior ectosylvian gyrus. **d, Combined verbal & somatosensory reinforcement** - 5s simultaneous reinforcement blocks where the examiner provided simultaneous praise (“Good girl Belka!”) & gently touching the left side of the face with eye contact, were interleaved with 5s rest periods. PSD from the anterior ectosylvian gyrus is shown. For analyses, PSDs for each task block were normalized by the average PSD over the whole experiment. Averaged normalized power was quantified for each frequency range in each task block. Task-associated changes were quantified using a signed r^2^ metric comparing active & inactive behavioral states(which can range from −1 to 1), as illustrated in the right-middle. All reported r^2^ are significant at p< 10^−5^ (unpaired t-test, Bonferroni corrected for number of channels). Note that data are common average re-referenced for generation of these PSDs (see Extended Data Fig. 1).

A researcher with an affectionate relationship with Belka performed five different sensory tasks in her familiar play area. Quietly petting the left side of the dog’s body activates the postcruciate gyrus (Figs. 2b&3), which evoked potential and fMRI studies have identified as the primary somatosensory region^11,12^. These previous studies also describe a second somatosensory area in the anterior ectosylvian gyrus (also called ectosylvia accessoria^13^), and we similarly found it to be active during quiet petting (Fig. 3). Surprisingly, this very same region was active if the dog was verbally praised from a distance without any physical touch (Figs. 2c&3), but was not active if a pure tone was played (Extended Data Fig. 2). Neural activity was maximal in these anterior ectosylvian gyrus measurements when petting and verbal praise were combined (Figs. 2d&3).

**Figure 3.**
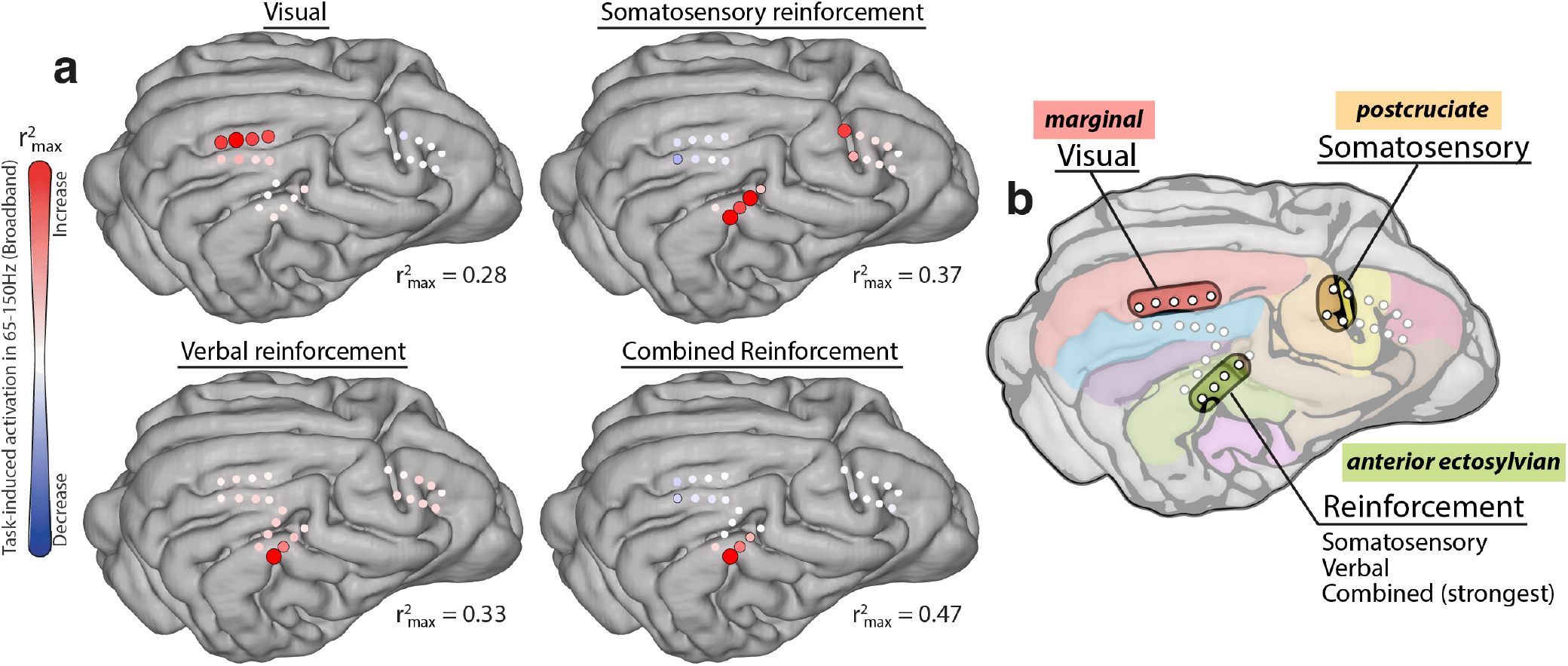
Functional maps across across sensory tasks. **a**, Maps of function in the canine brain showing distinct representation during each of four tasks. The electrode diameter and color intensity reflect task-associated revealed by broadband power changes in the PSD (65-150 Hz), quantified by r^2^ value (as in Fig. 2). Each task map is independently scaled for each task to 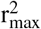, which is noted to the bottom right of each brain. A black circle around the channel denotes statistical significance at p< 0.05 by unpaired t-test, after Bonferroni correction for number of channels. Data were bipolar re-referenced for these maps (see Extended Data Fig. 1). Note that processing of a simple tone did not activate any area covered by the 3 grids (not shown here, see Extended Data Fig. 2). **b**, Summary plot showing functional representation. Note that the anterior ectosylvian gyrus, which is the canine anatomical homolog of human temporoparietal junction (Wernicke’s area), is active for social reinforcement, whether it is verbal or somatosensory (petting), and is most active for both combined.

What is the significance of these measurements? Independent higher-order somatosensory^11^ and auditory^14^ processing have suggested that the anterior ectosylvian gyrus is a site of multisensory integration, and our measurements corroborate this. Furthermore, auditory fMRI studies with canines have suggested that the social valence (by intonation of speech) can selectively augment brain activity throughout the ectosylvian gyrus^15^. Our combined observations from the implanted ECoG array suggest a deeper purpose to the anterior ectosylvian gyrus: Because it is not active when listening to a simple tone, but is active during quiet petting or verbal praise from a distance, and is even more active during simultaneous praise and petting, our data show that it is **an area that is specialized for social reinforcement** (Fig. 3, Extended Data Fig. 3).

Social reinforcement is of central importance to the lives of mammals (particularly those that live in groups), and evidence of specialized brain function in sensory processing of socially-reinforcing stimuli has been found across species. These have been particularly well established in basal ganglial and brainstem structures^16,17^. From a cortical perspective, the importance of processing socially-important sensory information is also found in many mammalian species^18^. For example, rodents have specialized circuitry in their auditory cortex for processing species-specific ultrasonic vocalizations that encode social reward and learning related to maternal care, mate choice, and other behavioral processes^19,20^. During social touch, single cells in rodent somatosensory cortex are modulated by the gender & mate context, independent of the physical properties of the touch itself^21^. However, the canine anterior ectosylvian gyrus shows something different and novel: a region that actively processes social reinforcement as an independent phenomenon, whether it arrives in a purely auditory or purely tactile fashion (Fig. 3).

While engagement of this region may have served as a substrate for canine domestication and cross-species social reinforcement, can this finding teach us something about our own brains? The last common ancestor of dogs and humans lived in the Cretaceous era ∼100 million years ago^22,23^. While we do not have biological samples of this ancestor that might inform direct anatomic homology, we do know from casts of the inner table of fossil skulls that no simple common brain sulcus motifs are preserved within the canine^24^ or primate^25^ evolutionary lineages. Instead, cross-species homology can be inferred from behaviorally-determined commonalities in functional representation of the cortex, and domains of projections from the thalamus and brainstem, which are evolutionarily better-preserved than cortex^26^. Canine primary auditory cortex has been localized to the middle and posterior portions of the ectosylvian gyrus based upon cortical lesioning^27^, tracing to the medial geniculate nucleus of the thalamus^28^, and fMRI^29^. The anterior ectosylvian hub of visual and auditory multisensory integration for social reinforcement that we find lies immediately rostral to the canine primary auditory cortex homolog. It has a unique pattern of connectivity, distinct from the auditory areas behind it^30^. In humans, the area immediately rostral to the primary auditory cortex, from an embyrological perspective, would lie at the transitional zone between the superior temporal gyrus and the supramarginal gyrus^31,32^ (the temporoparietal junction, Extended Data Fig. 4).

On both sides of the human brain, this cortical area is an important place for processing complex social reinforcement. Wernicke famously found that, in the dominant hemisphere (typically left) it is associated with receptive speech - how we understand what others want us to know^33^. More recent studies have found that, in the non-dominant hemisphere, this region is associated with theory of mind - how we model the internal thoughts and intentions of others^34^. Our finding of a homologous region in the dog suggests that the complex human functions of speech understanding and theory of mind may have evolved from a more general mammalian cortical structure that is specialized for social reinforcement.

## Methodology

### Neuromodulation platform

Our neuromodulation platform represents a novel fusion of an implanted device & a software ecosystem^4^. The implanted hardware consists of a CorTec BrainInterchange 32-channel sensing and stimulation device (BIC device), coupled to three small AirRay electrocorticography arrays (Fig. 1)^35^. It is powered by an inductively-coupled external source. BIC-specific modules within the BCI2000 software environment interfaces enable stimulation and recording^4,36^, and interface with the device by Bluetooth. This BrainInterchange-BCI2000 ecosystem is undergoing continuous development and optimization^4^. All software, data, resources, & protocols for this initiative are fully open-source, with a vast documentation to teach the community how to use it without having extensive technical expertise. A series of video tutorials and demonstrations for the platform has been created and is available at https://www.bci2000.org/mediawiki/index.php/CortecExperience. When fully developed, this ecosystem may enable clinical teams to create personalized closed-loop therapies tailored specifically to the needs of their patient population.

### Surgical implant

A 2-year-old female beagle, “Belka”, was implanted with the 32-channel sensing-and-stimulation Cortec BrainInterchange device as previously described^4^, according to a publicly available operative protocol^37^. Three arrays (32 electrocorticography channels) were implanted epidurally over the right hemispheric convexity with the FDA-approved AirRay electrodes^35^ (Extended Data Fig. 5). Sensory experiments were conducted at 11-13 months post-implant.

### Anatomic co-registration

A pre-implantation 3T MRI and a post-implantation CT were obtained. The brain was manually segmented from the MRI using 3D Slicer^38^, and the CT was co-registered with electrodes aligned to the anatomy using the CTMR package, as previously described^39^, which was also used for subsequent plotting. Anatomic segmentation of the brain surface was determined manually with reference to the *Stereotactic Cortical Atlas of the Domestic Canine Brain*^13^ (Extended Data Fig. 6). The 3 grids were localized to the 1) *frontal, precruciate* and *postcruciate* gyrus; 2) *ectosylvian gyrus* extending to the border of the *suprasylvian & ectomarginal* gyrus and the *rostral composite*; and 3) flanking the *marginal* and *ectomarginal* gyri. When bipolar re-referenced, analyzed data are plotted on the brain surface at the interpolated points between the bipolar pair.

### Canine tasks

Five types of tasks were performed - simple visual, simple tone, isolated somatosensory reinforcement, isolated verbal reinforcement, and combined somatosensory & verbal reinforcement. For each task, the four best runs (as determined by behavior, prior to data analysis) were selected for further analysis. Each run consisted of 15 repetitions of active & inactive task blocks.

#### Simple Visual (Fig. 2A)

Examination took place in a closed room with all external sources of light blocked. The dog was placed on a leash in the middle of the room, allowing her to move within the reach of the leash. 5-second “lights OFF” blocks with the room lights turned off were interleaved with 5s “lights ON” blocks, where the rooms lights were turned on. A laptop was in the room, with monitor light exposed, cuing the examiner to turn the room lights on and off.

#### Simple Tone (Extended Data Fig. 2)

Examination took place in a lit, closed, quiet room. The dog was sitting in the middle of the room, without feedback from the examiner. 5-seconds of a pure “A” tone (440 Hz) blocks were interleaved with 5s “quiet” blocks. A laptop was in the room, with the monitor exposed during the experiment.

#### Isolated Somatosensory Reinforcement (Fig. 2B)

The dog was positioned unrestrained within the examination room with the room lights on, next to the examiner. 3-second blocks of tactile stimulation (petting along the left side of the dog, encompassing the whiskers, front and hind limbs, and torso) were interleaved with 5s blocks of rest (Fig.2A). The examiner did not speak at all during the experiment.

#### Isolated Verbal Reinforcement (Fig. 2C)

The dog was positioned unrestrained within the examination room with the room lights on, next to, but not touching, the examiner. 3-second blocks of verbal reinforcement, saying “Good girl Belka” once each block, were interleaved with 5-second quiet blocks. No other reinforcement was given.

#### Combined Somatosensory & Verbal Reinforcement (Fig. 2D)

The dog was positioned unrestrained in the examination room next to the examiner with the room lights on and calming classical music playing at a low volume. 5-second reinforcement blocks where the examiner provided verbal reinforcement (“Good girl Belka!”) & gently touching the left side of the face with eye contact were interleaved with 5s rest periods.

### Electrophysiological measurements

Data were measured using BCI2000 general-purpose software^36^, which provides a graphical user interface for data acquisition, online processing for closed-loop application (though not used in this study), and stimulus presentation. Data were sampled at 1000 Hz, with an amplification gain of 57.5 dB / 1 *μ*V, and initially referenced to the channel 1 (Extended Data Fig. 1). Missing samples due to the packet loss were replaced with the first valid sample preceding the packet loss (see *Ayyoubi et al*. ^40^ for further packet loss discussion).

### Signal processing

Data were examined by raw trace as well as relative signal power to identify bad channels, which were discarded prior to either common-average or bipolar re-referencing of the data. Use of both of these referencing approaches is important because bipolar re-referencing suppresses coherent oscillations/rhythms while common average re-referencing falsely introduces oscillations into channels where they are otherwise absent, as illustrated in Extended Data Fig. 1. Power spectral densities (PSDs) up to 150Hz were calculated for each task block using Welch’s averaged periodogram method^41^ with 1s Hann windowing^42^ and 50% window overlap. Several data blocks were rejected (across all channels) due to significant transient artifact. Individual block PSDs were normalized by the mean power at each frequency (mean calculated over each full task). Signed *r*^*2*^ cross-correlations comparing task conditions (Fig. 2C) were calculated across all the channels at each frequency, and plotted on featuremaps (Extended Data Figs. 1-3) to characterize the spatial and frequency-specific structure of neurophysiological changes associated with each task. Separately, a high-frequency broadband range was chosen *a priori* at 65-150Hz to capture the 1/f structure that has been shown in humans to be a correlate of local population activity^5,6^. Averaged power (after normalizing by the mean PSD) was calculated across each frequency range for each block. Blocks of each type within each task were then compared with one another using a signed r^2^ metric and an unpaired (2-sample) t-test (mean normalized power in band for trials of petting vs. rest, lights on vs. lights off, tone vs. quiet, speech vs. quiet, and combined reinforcement vs. rest). Maps of r^2^ values were projected onto the rendered brain to show task-associated brain activity (Fig. 3), and channels that reached threshold significance (p<0.05 after Bonferroni correction), were marked with a black outline.

### Ethics statement

This research is conducted under Mayo Clinic IACUC protocol A00001713-16-R19. We maintain our canines in an IACUC-approved environment. In addition, according to State of Minnesota statute 135A.191, the canines will be made available for adoption at the conclusion of research. In the event of serious illness or decline, the animals may be humanely euthanized by the veterinary team according to an IACUC mandated protocol. The canine subject, Belka, is a 3 year old female (implanted at 2 years old). She is housed in a communal environment, and receives daily social interactions with veterinary staff as well as open time with other canines. The intent of this animal research is to test and develop a platform for novel human therapeutics, that may also be implanted into public pet dogs to treat their epilepsy (as we have done previously)^43^.

## Data and software availability

All recorded data and MATLAB analysis code available at [pointer to data archive here on final version].

## Manuscript preparation and AI utilization

Line art illustration panels in Fig. 2 were made with assistance of DALL-E 3 and subsequently modified by hand.

## Acknowledgements

We are grateful to be able to work with this animal, Belka, and for the care and attention provided by the Veterinary staff at Mayo Clinic. Belka will be available for adoption at the conclusion of the research in accordance with the Minnesota state Beagle Freedom Bill. This work was supported by the NIH U01-NS128612 (KJM, GAW, PB). The contents of this manuscript are solely the responsibility of the authors and do not necessarily represent the official views of the NIH. KJM was also supported by the Foundation For OCD Research. Our funders played no role in data collection and analysis, study design, decision to publish, or manuscript preparation.

## Author contributions

The research was conceived by KJM, GS, NI, PB, and GAW. KJM and IK performed the surgery, helped by pre- and intra-operative consultation with VAC. KJM, MAVdB, IK, and DH performed targeting, image segmentation and rendering. Animal maintenance, diagnostic measurement, and daily interactions were performed by FL, FM, JK, MAVdB, VK, and GAW. Behavioral sensory tasks were performed by FL. KJM and FL performed signal analyses. KJM drafted the manuscript with editing assistance by MRB, FL, and DH.

---

**Extended Data Fig. 1.**
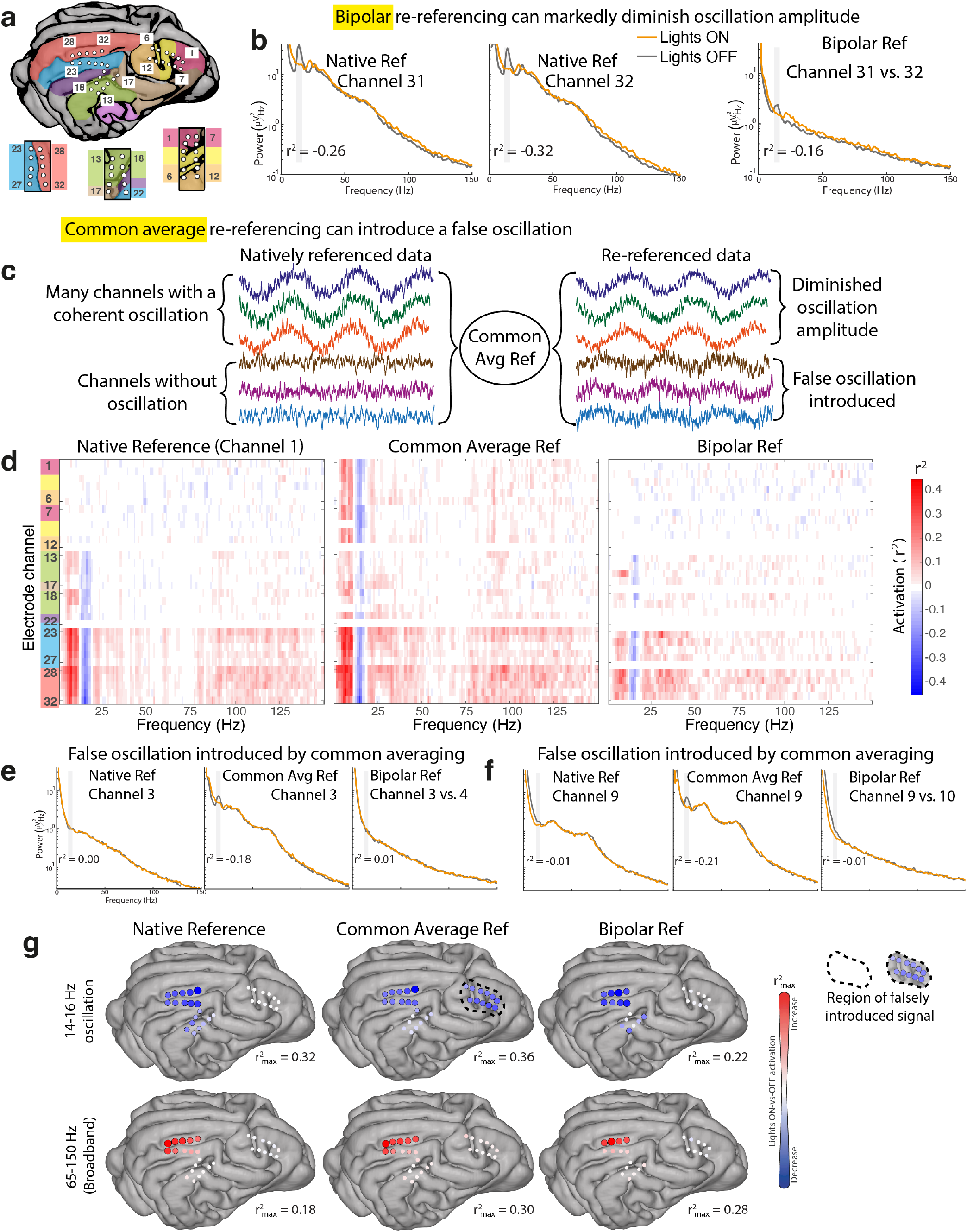
The effect of referencing on signal, illustrated for the lights-on vs. lights-off visual task. **a**, Electrode channel positions show locations of recording. **b**, Power spectral densities from channels 31 & 32 show an example of an ∼15Hz oscillation that is clearly present in the natively-referenced (to channel #1) during the lights-off condition. **c**, This example with synthetic data illustrates the effect of common-average re-referencing in an array with an oscillation that is widespread across many channels and coherent. Firstly, the oscillation will be diminished in the channels where it is natively present, and, secondly, a false oscillation will be introduced (180° phase-shifted). **d**, Featuremaps show r^2^ values in 1Hz bins as a function of frequency and channel comparing lights-on vs lights-off conditions (Fig. 2), reflecting state discriminability (“Activation”) in a task. The global effect of reference on discriminability is seen by comparing 3 regimes: natively-referenced data to channel 1 (left), common-average re-referencing (middle), and bipolar re-referencing (right). **e&f**, Illustrations for how common-averaging can falsely introduce an oscillation for channels 3 and 9. **g**, Brain renderings showing the effect of reference spatially on an oscillation (upper row) and broadband change (lower row). Black circle indicates p< 0.05 by unpaired t-test, after Bonferroni correction for number of channels.

**Extended Data Fig. 2.**
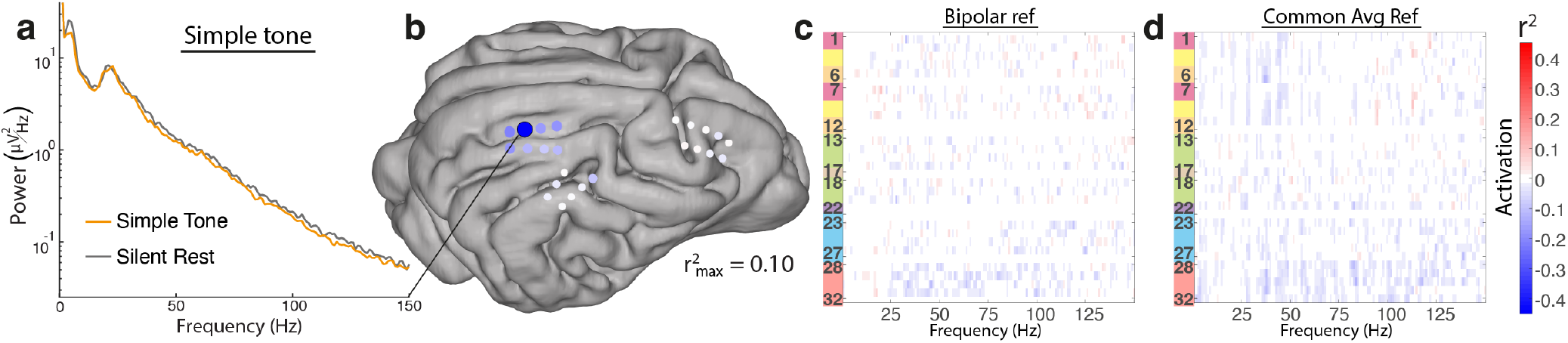
Brain functional activation map for a Simple tone task, where 5-seconds audio of a pure “A” tone (440 Hz) blocks were interleaved with 5s quiet blocks. **a**,**b**, PSD and brain map of pure tone during rest from the only channel that showed significant broadband change - a suppression during playing of the tone (bipolar reference). **c**,**d**, Featuremaps of r^2^ values from bipolar and common-average re-referenced data showing a paucity of task-associated change when playing the pure “A” tone.

**Extended Data Fig. 3.**
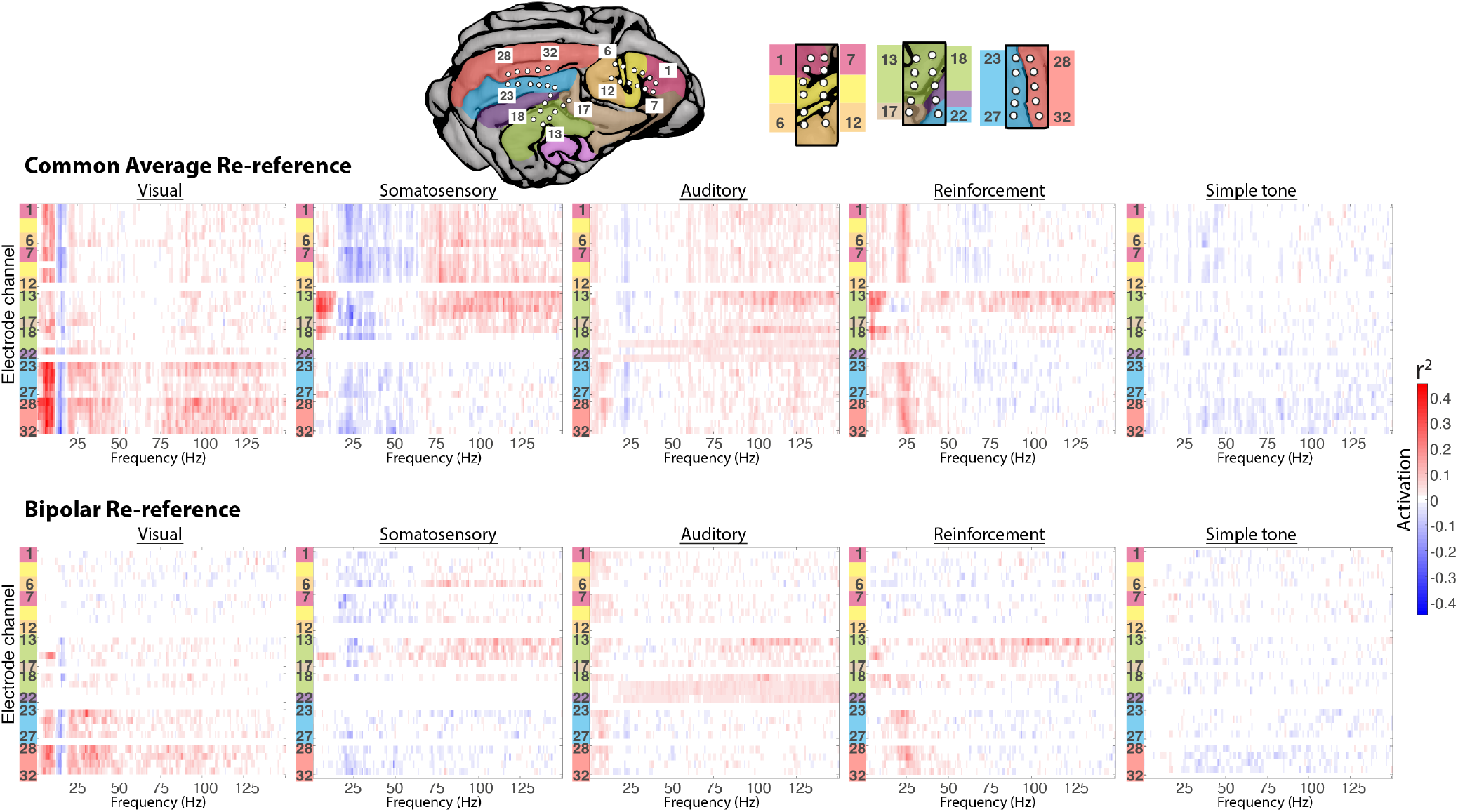
Featuremaps for all tasks. The top row of r^2^ featuremaps is for common-average re-referenced data, and the bottom row is for bipolar re-referenced data (all to the same scale). Except for the *Simple tone* task, each of the tasks is illustrated in Fig. 2.

**Extended Data Fig. 4.**
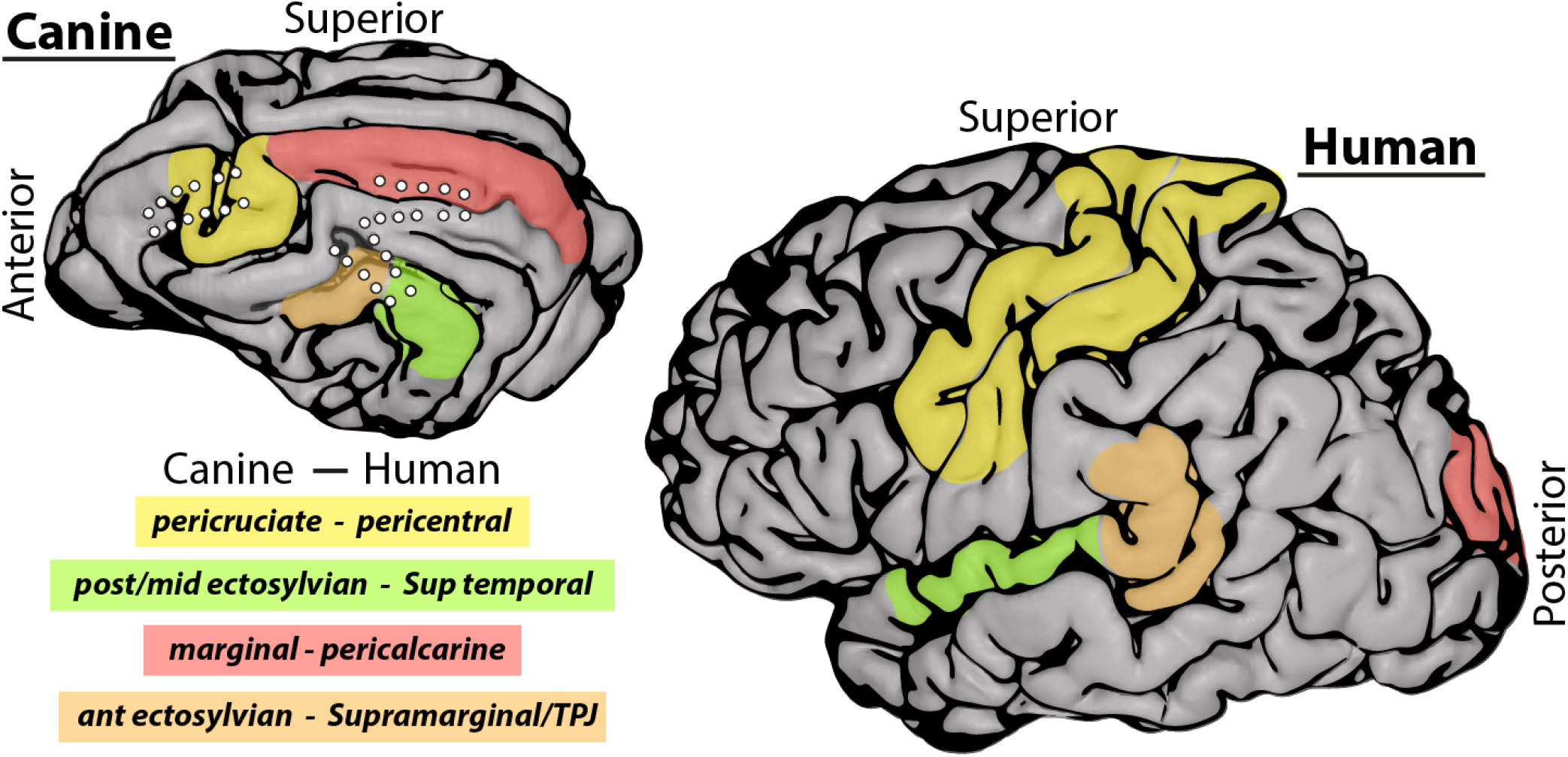
Homologous human and canine anatomic regions. Note that, while it may be appealing to assign anatomic homology to brain regions by the appearance of sulci, these similarities do not reflect evolutionarily preserved folds^24,25^. Instead, homology can be inferred from behaviorally-determined commonalities in functional representation of the cortex and domains of projections from the thalamus and brainstem, which are evolutionarily better-preserved than cortex^26^. These comparisons have previously identified homologous primary sensorimotor (**yellow shading**), auditory (**green shading**), and visual (**red shading**) regions. This hub of visual and auditory multisensory integration for social reinforcement that we find in this article in the anterior ectosylvian gyrus of the canine (**orange shading**) lies immediately rostral to the canine primary auditory cortex homolog^27,28^. In humans, the area immediately rostral to the primary auditory cortex, from an embyrological perspective, would lie at the transitional zone between the superior temporal gyrus and the supramarginal gyrus^31^ (temporoparietal junction - TPJ, **orange shading**).

**Extended Data Fig. 5.**
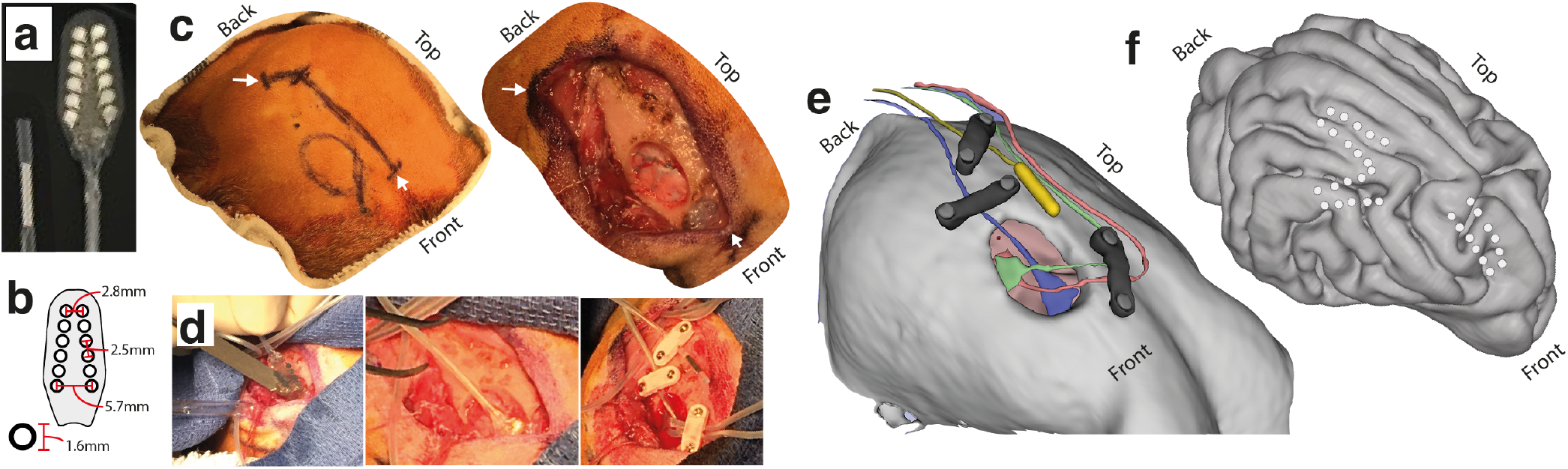
Electrode array surgical implantation. **a**, The ground electrode and one of the 2×6 angled electrode arrays used for recording. **b**, Schematic with dimensions of electrode array. **c**, Planned incision and craniotomy on scalp (left) with opening and craniotomy, showing epidural space (right). White arrows correspond to same location on left and right. **d**, Insertion of arrays into the epidural space and anchoring of electrodes. **e**, Rendering of skull and craniotomy showing electrode arrays in situ, and the ground electrode in yellow. Plates that hold lead in place are shown in black. **f**, Brain rendering showing three arrays in situ, extracted from pre-implant MRI and post-implant CT. *Note that the final two electrodes from the middle and posterior arrays were excluded, leaving a 5×2 configuration for recording*.

**Extended Data Fig. 6.**
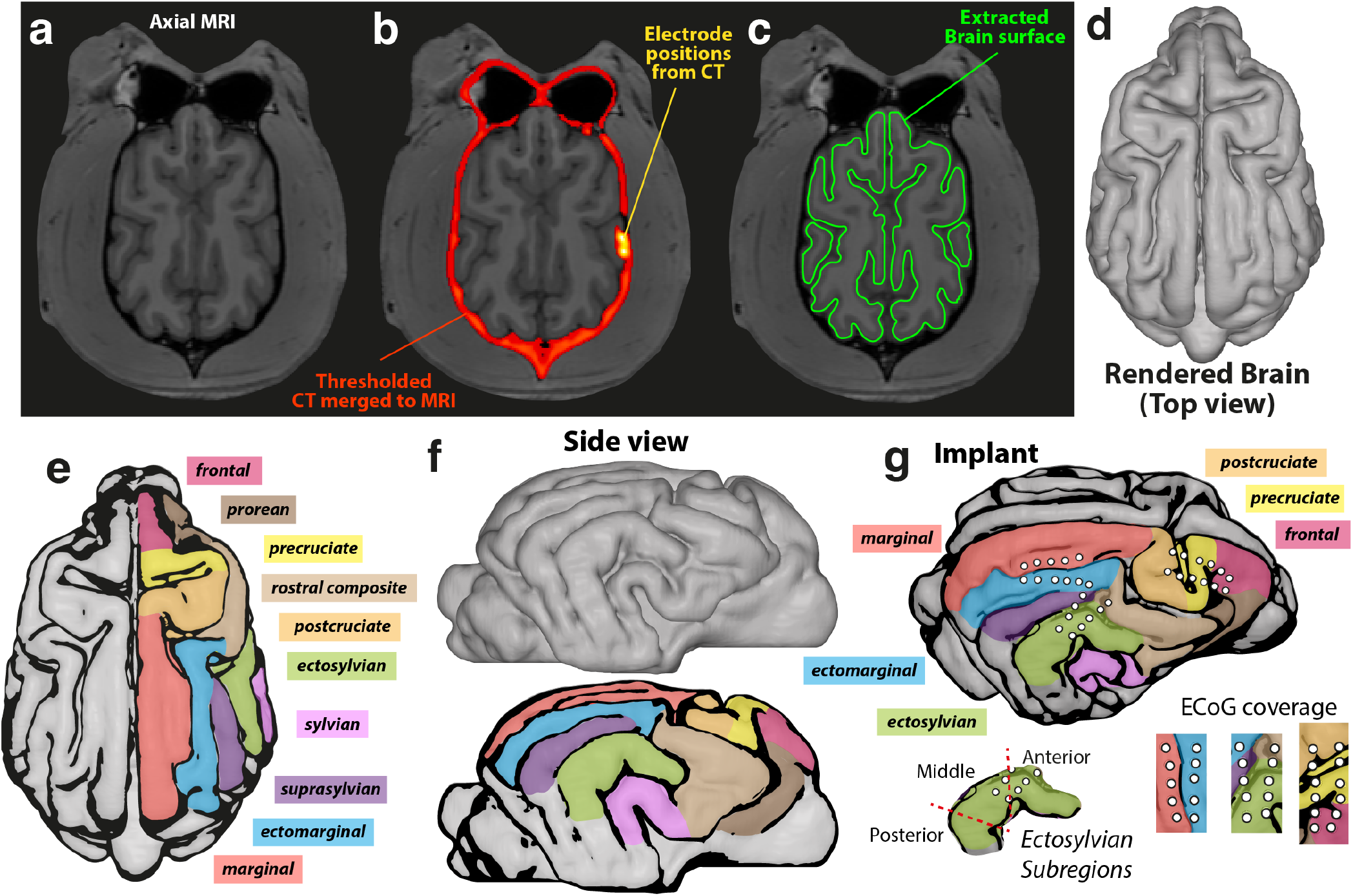
Segmentation of the canine brain surface from MRI and co-registration of electrode position to brain anatomy. **a**, A pre-operative volumetric T1 MRI was obtained with the canine fully anesthetized. **b**, A post-operative volumetric CT scan is co-registered to the MRI in order to identify ECoG electrode positions in relation to brain anatomy. **c**, The brain surface is segmented manually on cross sections of the MRI. **d**, A volumetric rendering of the brain surface is generated from the manual segmentation. **e**, Gyral anatomy is compared by direct reference to atlas anatomy^13^. **f**, A side view (from the right) of the brain rendering and gyral segmentation. **g**, The unified segmentation, gyral labeling, and co-registered ECoG electrode positions.

